# Genome-wide association study for maize leaf cuticular conductance identifies candidate genes involved in the regulation of cuticle development

**DOI:** 10.1101/835892

**Authors:** Meng Lin, Susanne Matschi, Miguel Vasquez, James Chamness, Nicholas Kaczmar, Matheus Baseggio, Michael Miller, Ethan L. Stewart, Pengfei Qiao, Michael J. Scanlon, Isabel Molina, Laurie G. Smith, Michael A. Gore

## Abstract

The cuticle, a hydrophobic layer of cutin and waxes synthesized by plant epidermal cells, is the major barrier to water loss when stomata are closed at night and under water-limited conditions. Elucidating the genetic architecture of natural variation for leaf cuticular conductance (*g*_c_) is important for identifying genes relevant to improving crop productivity in drought-prone environments. To this end, we conducted a genome-wide association study of *g*_c_ of adult leaves in a maize inbred association panel that was evaluated in four environments (Maricopa, AZ, and San Diego, CA in 2016 and 2017). Five genomic regions significantly associated with *g*_c_ were resolved to seven plausible candidate genes (ISTL1, two SEC14 homologs, cyclase-associated protein, a CER7 homolog, GDSL lipase, and β-D-XYLOSIDASE 4). These candidates are potentially involved in cuticle biosynthesis, trafficking and deposition of cuticle lipids, cutin polymerization, and cell wall modification. Laser microdissection RNA sequencing revealed that all these candidate genes, with the exception of the CER7 homolog, were expressed in the zone of the expanding adult maize leaf where cuticle maturation occurs. With direct application to genetic improvement, moderately high average predictive abilities were observed for whole-genome prediction of *g*_c_ in locations (0.46 and 0.45) and across all environments (0.52). The findings of this study provide novel insights into the genetic control of *g*_c_ and have the potential to help breeders more effectively develop drought-tolerant maize for target environments.

**Article summary:** The cuticle serves as the major barrier to water loss when stomata are closed at night and under water-limited conditions and potentially relevant to drought tolerance in crops. We performed a genome-wide association study to elucidate the genetic architecture of natural variation for maize leaf cuticular conductance. We identified epidermally expressed candidate genes that are potentially involved in cuticle biosynthesis, trafficking and deposition, cutin polymerization, and cell wall modification. Finally, we observed moderately high predictive abilities for whole-genome prediction of leaf cuticular conductance. Collectively, these findings may help breeders more effectively develop drought-tolerant maize.

## Introduction

The cuticle is a hydrophobic layer covering the surface of the shoot, which limits transpiration, in addition to protecting shoot tissues from UV radiation, heat, and pathogen attack (Shepherd and Wynne Griffiths 2006; Xue *et al*. 2017). Cuticles have two major lipid components: cutin and waxes. Cutin is a polymer composed of fatty acid derivatives and glycerol, and is highly cross-linked to form an insoluble matrix (Pollard *et al*. 2008; Fich *et al*. 2016). Soluble waxes are deposited into and on top of this matrix and consist mainly of aliphatic compounds derived from very-long-chain fatty acids including alkanes, aldehydes, alcohols, ketones, and wax esters (Yeats and Rose 2013). The major pathways for both wax and cutin monomer biosynthesis have been elucidated via genetic and biochemical studies conducted mainly on model plant systems (Yeats and Rose 2013; Lee and Suh 2015; Fich *et al*. 2016). Many transcriptional regulators of cuticle biosynthesis have also been identified (Borisjuk *et al*. 2014). Pathways for delivery of cuticle lipids from the intracellular membranes where they are synthesized to the extracellular environment, and to their final destinations external to the cell wall, have been partially elucidated. Golgi-mediated vesicle trafficking (McFarlane *et al*. 2014), ATP BINDING CASSETTE TRANSPORTER G (ABCG) family proteins (Pighin *et al*. 2004; Bird *et al*. 2007; Panikashvili *et al*. 2010; Bessire *et al*. 2011; Chen *et al*. 2011), and extracellular lipid transfer proteins (DeBono *et al*. 2009; Kim *et al*. 2012) are required for accumulation of multiple classes of cuticle lipids at the shoot surface, but how these transport processes work together is not well understood.

Cuticle composition and structure varies widely among species and tissue types (Jetter *et al*. 2008), and its permeability to water can vary as much as three orders of magnitude (Kerstiens 2006). Relationships between cuticle composition, structure, and water barrier function are complex. The simple idea that cuticle impermeability to water increases with increased wax load or cuticle thickness has been repeatedly refuted; instead, cuticle composition and the organization of components appear to determine water barrier function (Riederer and Schreiber 2001). Cuticle composition is also modulated by environmental factors. Light, temperature, and osmotic stress influence both the quantity and composition of cuticular waxes (Shepherd and Wynne Griffiths 2006; Kosma and Jenks 2007). While high relative humidity usually suppresses wax production (Sutter and Langhans 1982; Koch *et al*. 2006), drought stress has been found to change cuticular wax composition (Panikashvili *et al*. 2007; Kosma *et al*. 2009) and increase wax deposition in several crop species (Shepherd and Wynne Griffiths 2006; Cameron *et al*. 2006; Kosma and Jenks 2007).

In addition to modulation of its composition by drought stress, a variety of other observations support a role for the cuticle in drought tolerance. Most mutations and other genetic modifications affecting cuticle composition or wax load increase its permeability to water, and this has often been associated with decreased drought tolerance (Zhou *et al*. 2013; Zhu and Xiong 2013; Li *et al*. 2019). In a few cases, overexpression of cuticle lipid biosynthetic enzymes, or their transcriptional regulators, increased cuticular lipid abundance and drought tolerance (Aharoni *et al*. 2004; Zhang *et al*. 2005; Bourdenx *et al*. 2011; Wang *et al*. 2012). Glaucousness (a visible trait resulting from abundant accumulation of epicuticular wax crystals on shoot tissue surfaces) was selected for during the domestication of wheat and involves genes regulating wax biosynthesis (Hen-Avivi *et al*. 2016; Bi *et al*. 2016); analyses of heritable variation in glaucousness in wheat and barley has revealed positive correlations between this trait and drought tolerance (Febrero *et al*. 1998; Guo *et al*. 2016). These findings point to the potential relevance of cuticle modification for increasing drought tolerance in cereal crops.

There is a longstanding interest in breeding for cuticle-related traits such as drought tolerance, but the complexity of the relationship between cuticle composition and resistance to dehydration, and the lack of methods amenable to high-throughput phenotyping for cuticle-associated traits, has hampered progress in this area (Petit *et al*. 2017). Genome-wide association studies (GWAS) (Yu *et al*. 2006; Zhang *et al*. 2010; Lipka *et al*. 2015), taking advantage of natural variation in cuticle permeability and historical recombination events within a single species, offer an attractive approach to identifying genes and alleles that can decrease cuticle permeability. Additionally, findings from GWAS of cuticle-related phenotypes could be used to better inform the application of genomic selection strategies (Meuwissen *et al*. 2001; Lorenz *et al*. 2011) for increasing drought tolerance in crop species.

In this study, we utilized GWAS combined with laser microdissection RNA sequencing (LM-RNAseq) of epidermal tissue samples, from zones of the expanding maize leaf where the cuticle develops, to implicate candidate genes controlling the water barrier function of the maize leaf cuticle. Methods directly measuring permeability of the cuticle to water (Valeska Zeisler-Diehl *et al*. 2017) are not amenable to the high-throughput phenotyping needed for GWAS involving analysis of thousands of samples. Thus, we utilized a phenotyping strategy that indirectly measures cuticle water barrier function by calculating the drying rates of detached leaves placed in the dark to close stomata (Ristic and Jenks 2002). While some water may be lost via stomata that are not completely sealed, the sealing of stomata is thought to depend on cuticular flaps lining the stomatal pore that overlap when stomata are closed (Zhao and Sack 1999; Kosma and Jenks 2007). Thus, we used this phenotyping strategy with the expectation that it primarily measures cuticle-dependent water loss, which we refer to as adult leaf cuticular conductance (*g*_c_). Maize is most sensitive to drought stress at flowering (Grant *et al*. 1989), when juvenile leaves have already senesced and only adult leaves remain. Thus, we utilized adult leaves of field-grown plants for this analysis. Moreover, since cuticle composition is known to vary depending on the growth environment (Shepherd and Wynne Griffiths 2006), we utilized plants grown in two contrasting environments: the cooler and more humid environment of San Diego, CA and the hotter and more arid environment of Maricopa, AZ. We aimed to (i) detect genomic regions associated with natural variation in *g*_c_; (ii) enhance selection of candidate genes impacting this trait through an LM-RNAseq analysis of the expanding maize adult leaf; and (ⅲ) evaluate whole-genome prediction models to facilitate genomic selection on this trait, providing tools for potential improvement of drought-tolerance in maize for target environments.

## Materials and Methods

### Plant materials and experimental design

A set of 468 maize inbred lines from the Wisconsin Diversity panel (Hansey *et al*. 2011) was evaluated for adult leaf cuticular conductance (*g*_c_). The inbred lines were planted at the Maricopa Agricultural Center, Maricopa, AZ, and University of California San Diego, San Diego, CA, in 2016 and 2017. The layout of the experiment in each of the four environments (location × year combination) was arranged as an 18 x 26 incomplete block design. Each incomplete block of 18 experimental lines was augmented by the random positioning of two checks (B73 and Mo17, 2016; N28Ht and Mo17, 2017). The entire experiment of 468 unique inbred lines and repeated checks was grown as a single replicate in each environment. Edge effects were reduced by planting a locally adapted maize inbred line around the perimeter of the experiment. Experimental units were one-row plots of 3.05 m (Maricopa) and 4.88 m (San Diego) in length with 1.016 m inter-row spacing. There was a 0.91 m alley at the end of each plot. In each plot, 12 kernels were planted, followed by thinning the stand as needed to limit plant-to-plant competition. We obtained and summarized meteorological data (Table S1) from automated weather stations located at a distance of ∼200 m or less from the experimental field sites in Maricopa, AZ (Arizona Meteorological Network; http://ag.arizona.edu/azmet/index.html) and San Diego, CA (Scripps Hydroclimate Weather Station).

### *g*_c_ evaluation and phenotypic data analysis

Initially, a porometer (model SG-1, Decagon Devices, Pullman, WA) was used to measure diffusion conductance for adult leaves collected from the field in the morning (at the same time of day that leaves were harvested for *g*_c_ analysis, within a week of their anthesis dates), and transferred to a dark room where they were kept with cut leaf bases immersed in water throughout the analysis. Diffusion conductance was measured at a series of timepoints for a selection of lines that had previously been demonstrated to have high *g*_c_, and B73 as a reference standard harvested each sampling day. As shown in Figure S1, most lines reached minimum conductance values, indicating stomatal closure, within 30 min, and all lines reached minimum values within 90 min. Based on these findings, we concluded that 2 h in the dark was sufficient to achieve stomatal closure, even for lines with high *g*_c_.

The findings from the porometer study were used to inform our phenotyping of the Wisconsin Diversity panel. To inform sampling times and help control for maturity differences, flowering time (days to anthesis, DTA) was recorded as the number of days from planting to initiation of pollen shed for 50% of the plants in a plot. For each plot, *g*_c_ was evaluated on an adult leaf from five plants when 50% of the plants in the plot were at anthesis. From each of five plants, the leaf subtending the uppermost ear, or the leaf immediately above or below it, was excised at 2.5 cm below the ligule. If there were fewer than five plants to measure in a plot, two, three, or four plants were evaluated to represent the *g*_c_ of that plot. Cut ends were immersed in water and leaves were incubated in a dark, well-ventilated room for 2 h at 20-22 °C and 55-65% RH to close stomata while ensuring that the leaves were fully hydrated. Subsequently, each leaf was hung on a line from a boot clip in the same dark, temperature-and humidity-controlled room, after carefully wiping the leaves to remove excess water. In 2017, the cut leaf end was wrapped in parafilm when hung to further reduce the incidence of non-cuticular transpiration. The wet weight of each leaf was then recorded every 45 min over a time period of 3 h, for a total of five measurements per leaf. Leaves were subsequently dried at 60 °C for 4 d in a forced-air oven, and afterward weighed.

To investigate the degree to which dry weight approximates surface area, we constructed an imaging table that consisted of a digital camera (Canon PowerShot S110) mounted at a fixed distance and folding table top to help flatten the adult maize leaves. In 2017, each experimental leaf that had completed the *g*_c_ phenotyping process was flattened, imaged, and then oven-dried as earlier described and weighed. Each image contained 4 to 7 leaves, depending on their sizes. Next, images were individually analyzed with a custom ImageJ (Schneider *et al*. 2012) macro to estimate the surface area (mm^2^) of each leaf. Even though every attempt was made to flatten the leaves, there were frequent occurrences of where wavy leaves could not be completely flattened. Very strong Pearson’s correlations were observed between the plot-level averages of leaf dry weight and surface area in MA (*r* = 0.93) and SD (*r* = 0.92) in 2017 (Figure S2; Table S2). Given these strong correlations and challenges in measuring surface area, leaf dry weight was considered to be a reasonable approximation of leaf surface area, and was used in the calculation of cuticular conductance as described below.

Adult leaf cuticular conductance (*g*_c_) from unit surface area was calculated as follows:

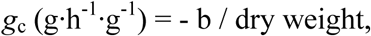

where b (g·h^-1^) is the coefficient of the linear regression of leaf wet weight (g) on time (h), and dry weight (g) is an approximation of leaf surface area.

To screen the phenotypic data (flowering time or *g*_c_) for significant outliers, univariate mixed linear models were fitted as follows: (1) each single environment; (2) the two environments in each location; and (3) all four environments. The model terms included grand mean and check as fixed effects and environment, genotype, genotype-by-environment (G×E) interaction (only for Models 2 and 3), incomplete block within environment, and column within environment as random effects. The Studentized deleted residuals (Neter *et al*. 1996) generated from these mixed linear models were assessed and significant (α = 0.05) outliers removed. For each outlier screened phenotype, an iterative mixed linear model fitting procedure was conducted for each of the three (1 to 3) full models in ASReml-R version 3.0 (Gilmour *et al*. 2009). All random terms that were not significant at α = 0.05 in a likelihood ratio test (Littell *et al*. 2006) were removed from the model, allowing a final best-fit model to be obtained for each phenotype. These final models (Table S3) were used to generate a best linear unbiased predictor (BLUP) for *g*_c_ and DTA for each line (Table S4).

Variance component estimates from the fitted full models (Table S5) were used to estimate heritability on a line-mean basis (Holland *et al*. 2010; Hung *et al*. 2012) for each phenotype in a location (Model 2) and across all four environments (Model 3). Standard errors of the heritability estimates were calculated with the delta method (Lynch *et al*. 1998; Holland *et al*. 2010). Pearson’s correlation coefficients were used to evaluate the strength of correlation between BLUP values for *g*_c_ and DTA in all pairwise combinations. The significance of correlations (α = 0.05) was tested using the function “cor.test” in R version 3.5.1 (R core team, 2018).

### DNA extraction and genotyping

For each inbred line, total genomic DNA was isolated from a leaf tissue sample consisting of bulked young leaves from a single plant. The leaf tissue samples were lyophilized and ground using a GenoGrinder (Spex SamplePrep, Metuchen, NJ, USA), followed by isolation of total genomic DNA using the DNeasy 96 Plant Kit (Qiagen Incorporated, Valencia, CA, USA). The genotyping-by-sequencing (GBS) procedure was conducted on the DNA samples following Elshire *et al*. (2011) with *Ape*KI as the restriction enzyme at the Cornell Biotechnology Resource Center (Cornell University, Ithaca, NY, USA). The constructed 192-plex GBS libraries were sequenced on a NextSeq 500 (Illumina Incorporated, San Diego, CA, USA).

We called genotypes at 955,690 high-confidence single-nucleotide polymorphism (SNP) loci in B73 RefGen_v2 coordinates following the method of Baseggio *et al*. (2019). Briefly, the raw SNP genotype calls were initially filtered to exclude singleton and doubleton SNPs (a minor allele observed in only a single line) and retain only biallelic SNPs having a SNP call rate greater than 10% and minimum inbreeding coefficient of 0.8 per Romay *et al*. (2013). Additionally, only lines with a call rate greater than 40% were retained. Missing SNP genotypes were partially imputed using FILLIN (Swarts *et al*. 2014) with a set of maize haplotype donors files having a 4 kb window size (AllZeaGBSv2.7impV5_AnonDonors4k.tar.gz, available at panzea.org). Given that missing genotype data still remained following partial imputation, the imputed genotype data set was further filtered to retain SNPs with a minimum call rate of 0.6, a minimum inbreeding coefficient of 0.8, and a minimum minor allele frequency of 5%. The final complete set contained 235,004 high-quality SNP markers scored on 451 lines that had a BLUP value for *g*_c_ in one or more environments. The genome coordinates of the SNP loci were uplifted by aligning 101 bp context sequences containing target SNPs (± 50 bp) to the B73 RefGen_AGPv4 reference genome with *Vmatch* (Kurtz 2003).

### Population structure analysis

We estimated population structure from the 451 line × 235,004 SNP genotype matrix in fastSTRUCTURE version 1.0 (Raj *et al*. 2014), with the number of ancestral populations (K) varying from 1 to 10 using the simple prior. To complement population structure inference with fastSTRUCTURE, a principal component analysis (PCA) was performed on the identical genotype matrix with the prcomp function in R version 3.5.1 (R core team, 2018). Next, results from fastSTRUCTURE and PCA were jointly interpreted and complemented with pedigree information (Hansey *et al*. 2011) to choose K=6 as the number of subpopulations. Additionally, lines were classified following group assignments of Hansey *et al*. (2011) (Figure S3A). In fastSTRUCTURE, the simple prior approach was used with K=6 to calculate the subpopulation composition of each line (Figure S3B). Lines with an assignment value of Q ≥ 0.50 were assigned to subpopulations (1, 2, 3, 4, 5, or 6), while those with assignment values of Q < 0.50 for all six subpopulations were classified as admixed (Figure S3C; Table S6). This information was used to inform a stratified sampling approach used for whole-genome prediction (WGP).

### Genome-wide association study

To identify associations between each of the 235,004 SNP markers and the BLUP values of *g*_c_ from the 451 inbreds, a GWAS was conducted with a univariate mixed linear model that employed population parameters previously determined (P3D) approximation (Zhang *et al*. 2010) in the R package Genome Association and Prediction Integrated Tool (GAPIT) version 3.0 (Lipka *et al*. 2012). To control for phenotypic variation due to maturity, population structure, and unequal relatedness, the fitted mixed linear models included BLUPs of flowering time (DTA) of corresponding environments, principal components (PCs), and a genomic relationship (kinship) matrix. The kinship matrix based on the centered IBS method (Endelman and Jannink 2012) implemented in TASSEL 5.0 (Bradbury *et al*. 2007) was calculated from a subset of 41,259 SNPs remaining after linkage disequilibrium (LD) pruning (*r^2^* ≤ 0.2) of the complete marker data set in PLINK version 1.09_beta5 (Purcell *et al*. 2007). The optimal number of covariates (BLUPs of DTA and PCs based on the genotypic matrix) to include in the mixed linear model was determined with the Bayesian information criterion (BIC) (Schwarz 1978). Prior to conducting the GWAS, the remaining missing genotypes for all SNP loci were conservatively imputed as a heterozygote in GAPIT. A likelihood-ratio-based *R*^2^ statistic (*R*^2^_LR_) (Sun *et al*. 2010) was used to estimate the amount of phenotypic variation explained by the model with or without a tested SNP. The false discovery rate (FDR) was controlled at 10% with the Benjamini-Hochberg procedure (Benjamini and Hochberg 1995).

### Linkage disequilibrium

The squared allele-frequency correlation (*r*^2^) method (Hill and Weir 1988) was used to estimate linkage disequilibrium between all pairs of SNPs on a chromosome in TASSEL 5.0 (Bradbury *et al*. 2007). The complete set of 235,004 high-quality SNPs was used for LD estimation; however, the missing SNP genotypes remaining after FILLIN imputation were not imputed with a heterozygote value prior to LD analysis. The estimates of LD were used to approximate the physical distance at which LD decayed to genome-wide background levels (*r^2^*< 0.10) and assess the pattern of LD surrounding GWAS-detected SNPs.

### Candidate gene identification

We searched for candidate genes within ±200 kb of the most significant SNP (peak SNP) for each detected genomic region associated with *g*_c_. Gene functional annotations (B73_RefGen_v4) of identified candidate genes were obtained from Gramene (http://www.gramene.org/). Additionally, the Gramene Mart tool was used to identify homologs of candidate genes in *Arabidopsis thaliana* (L.) Heynh. (Columbia-0 ecotype) and rice (*Oryza sativa* L. ssp. *Japonica* cv. ‘Nipponbare’). Gene functional annotations for the Arabidopsis and rice homologs were obtained from TAIR (https://www.arabidopsis.org/) and RAP-DB (https://rapdb.dna.affrc.go.jp/), respectively.

### Transcript abundance analysis for candidate genes

To investigate the patterns of transcript abundance for candidate genes potentially regulating *g*_c_ of maize leaves, we analyzed the LM-RNAseq dataset of Qiao et al. (2019) generated from epidermal and internal tissues that were LM from seven 2-cm-long intervals (six intervals from 2-14 cm, and one interval from 20-22 cm) from the expanding leaf 8 of maize inbred B73 with three biological replicates and three plants per replicate. The generated RNAseq reads were aligned to B73 genome RefGen_v4 with HiSAT2 (Kim *et al*. 2015) and counted with HTSeq (Anders *et al*. 2015). Transcript abundance levels for canonical protein coding genes were characterized as counts per million (cpm) after normalizing against library sizes with the R package edgeR 3.3.2 (Robinson *et al*. 2010). To identify potential outlier samples, genes with >1 cpm in two or more samples were retained. Next, a PCA was conducted on the cpm values of all the retained genes on all samples sequenced (Figure S4), and the epidermal and internal samples from interval 4-6 cm in the 3^rd^ replicate were removed as outliers (circled samples in Figure S4). With the remaining epidermis samples, a gene was declared expressed in the epidermis if its cpm was greater than one for at least three out of the 20 samples.

### Whole-genome prediction

WGP of *g*_c_ was conducted using a maximum likelihood approach to ridge regression in the ‘rrBLUP’ R package (Endelman 2011). The final complete set of 235,004 SNPs with post-FILLIN missing genotype data imputed as a heterozygote was used for WGP. We trained WGP models with *g*_c_ BLUP values from either MA (two environments), SD (two environments), or AllEnv (four environments), then each of the three models were separately used to predict *g*_c_ for MA, SD, and AllEnv. Collectively, this resulted in a total of nine prediction scenarios. Five-fold cross-validation as described in Owens *et al*. (2014) was performed to evaluate the predictive ability for *g*_c_ in each of the nine scenarios by calculating the Pearson’s correlation between observed BLUP values and genomic estimated breeding values. We used a stratified sampling approach to account for the presence of population structure, enabling each fold to have the proportion of subpopulations (1 to 6 and admixed) observed for the whole panel (Table S6). The procedure was conducted 50 times for each scenario, and the predictive ability was calculated as a mean of the correlations.

### Data availability

The raw GBS sequence data were deposited in the National Center of Biotechnology Information (NCBI) Sequence Read Archive (SRA) under accession number SRP160407 and in BioProject under accession PRJNA489924. The BLUP values of the phenotypes are provided in Table S4. The FILLIN partially imputed SNP genotype data for the 235,004 loci scored on the 451 inbred lines, weather data from Maricopa and San Diego in 2016 and 2017, and leaf image data from Maricopa and San Diego in 2017 are available at CyVerse (http://datacommons.cyverse.org/browse/iplant/home/shared/GoreLab/dataFromPubs/Lin_LeafCuticle_2019). The ImageJ macro for image analysis is available at Github (https://github.com/GoreLab/Maize_leaf_cuticle/blob/master/GWAS_CE/cliveMacro_2.4.txt). RNAseq reads of epidermal cells along the developmental gradient of the expanding leaf 8 of maize inbred B73 were deposited in the NCBI SRA under accession number SRP116320 and in BioProject under accession PRJNA400334, as described in Qiao et al. (2019).

Supplemental material available at FigShare: URL to be added

## Results

### Phenotypic variability

To establish the feasibility of GWAS for *g*_c_, the extent of phenotypic variation for adult leaf *g*_c_ was assessed in the Wisconsin Diversity panel of more than 450 maize inbred lines that was grown in two field locations (SD, San Diego, CA; and MA, Maricopa, AZ) in 2016 and 2017. The climatically contrasting environments of the SD and MA locations (Table S1) had the lowest and highest average *g*_c_ estimates, respectively (Table 1). There was a moderately strong correlation for *g*_c_ between years in a location (*r* = 0.52 and 0.54), whereas correlations were slightly weaker (*r* = 0.36 to 0.48) between locations within and across years (Figure S5). With respect to flowering time, *g*_c_ tended to have a moderately weak negative correlation with days to anthesis (DTA) in each environment (*r* = −0.14 to - 0.50), with the exception of a negative correlation of moderate strength (−0.50) in the MA17 environment (Figure S5). Comparatively, the first two principal components (PCs) but not the third PC calculated from the SNP genotype matrix (PC1, 3.68%; PC2, 1.83%; and PC3, 1.62%), which captures patterns of population structure, showed comparable correlations with *g*_c_ across all four environments (|*r*| = 0.14 to 0.31). Heritability on a line-mean basis across (AllEnv) locations was 0.71, with a range of 0.63 to 0.67 in locations (Table 1). These findings suggest that most phenotypic variation for *g*_c_ is under genetic control.

**Table 1.** Averages and ranges for best linear unbiased predictions (BLUPs) of adult maize leaf cuticular conductance (*g*_c_) in a location (MA and SD) and across all four environments (AllEnv), and estimated heritabilities on a line-mean basis.

| Environment | No. lines | BLUPs in $\text{g}\cdot\text{h}^{-1}\cdot\text{g}^{-1}$ | | | | | Heritability | |
| --- | --- | --- | --- | --- | --- | --- | --- | --- |
|  |  | Average | Min | Max | S.D.<sup>a</sup> | C.V.<sup>b</sup> | Estimate | S.E.<sup>c</sup> |
| MA | 410 | 0.20 | 0.15 | 0.29 | 0.03 | 0.15 | 0.63 | 0.04 |
| SD | 445 | 0.13 | 0.08 | 0.21 | 0.02 | 0.15 | 0.67 | 0.03 |
| AllEnv | 451 | 0.18 | 0.13 | 0.25 | 0.02 | 0.11 | 0.71 | 0.02 |

<sup>a</sup>Standard deviation of the BLUPs
<sup>b</sup>Coefficient of variation
<sup>c</sup>Standard error of the heritability

### Genome-wide association study

The genetic basis of natural variation for *g*_c_ was examined in the Wisconsin Diversity panel that had been genotyped with 235,004 high-quality SNP markers. We conducted GWAS in (MA or SD) and across locations (AllEnv) using a mixed linear model that controls for population structure and unequal relatedness. In addition, flowering time, which was also found to be moderately heritable (0.88 to 0.93) for MA, SD, and AllEnv (Table S7), was retained as a covariate for GWAS of *g*_c_ from MA and AllEnv based on the BIC values, thus attenuating the confounding effect of maturity when estimating allelic effects. A total of nine unique SNPs localized to five different genomic regions were significantly associated with *g*_c_ from MA and/or AllEnv at a genome-wide FDR of 10% (Figures 1 and S6). Of these nine SNPs, two were associated with *g*_c_ in both MA and AllEnv, for a total of 11 significant marker-trait associations (Table 2). No significant associations were detected for SNP markers with *g*_c_ from SD at the 10% FDR level (Figure S6).

**Figure 1.**
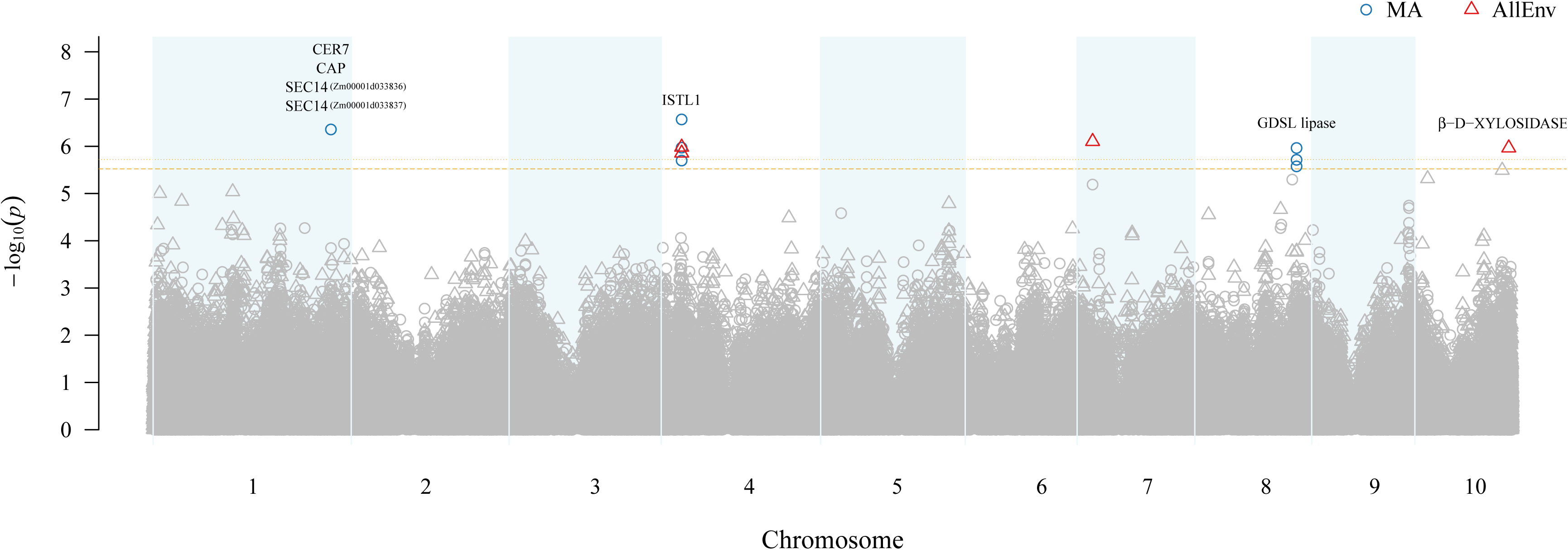
Manhattan plot of results from a genome-wide association study (GWAS) of adult maize leaf cuticular conductance (*g*_c_) conducted in Maricopa (MA) and across all four environments (AllEnv). The −log_10_ *P*-value of each SNP tested in a mixed linear model analysis of *g*_c_ is plotted as a point against its physical position (B73 RefGen_v4) for the 10 chromosomes of maize. The least significant single-nucleotide polymorphism (SNP) at a genome-wide false discovery rate of 10% in MA and AllEnv is indicated by a dashed horizontal orange line and a dotted horizontal orange line, respectively. SNPs significantly associated with *g*_c_ in MA and AllEnv are represented by blue circles and red triangles, respectively. The most plausible candidate genes within ± 200 kb of the significantly associated SNPs are listed above their corresponding GWAS signal.

**Table 2.** Significant SNPs detected at false discovery rate (FDR) of 10% through a genome-wide association study of adult maize leaf cuticular conductance (*g*_c_) in a location (Maricopa, MA; San Diego, SD) and across all four environments (AllEnv).

| SNP ID<sup>a</sup> | Chr | SNP position (bp)<sup>b</sup> | P-value | FDR-adjusted P-value<sup>c</sup> | Minor allele frequency | Allelic effect estimate<sup>d</sup> | $R^2_{LR}$<sup>e</sup> | $R^2_{LR-SNP}$<sup>f</sup> | Environments |
| --- | --- | --- | --- | --- | --- | --- | --- | --- | --- |
| S1_270551534 | 1 | 275,318,146 | 4.37E-07 | 0.05 | 0.13 | 0.009 | 0.28 | 0.33 | MA |
| S4_30201406 | 4 | 31,733,780 | 1.39E-06 | 0.08 | 0.07 | 0.009 | 0.30 | 0.34 | AllEnv |
|  | 4 | 31,733,780 | 1.06E-06 | 0.06 | 0.07 | 0.011 | 0.28 | 0.33 | MA |
| S4_30201436 | 4 | 31,733,810 | 2.68E-07 | 0.05 | 0.12 | 0.010 | 0.28 | 0.33 | MA |
| S4_30201599 | 4 | 31,733,973 | 1.04E-06 | 0.08 | 0.13 | 0.007 | 0.30 | 0.34 | AllEnv |
|  | 4 | 31,733,973 | 1.99E-06 | 0.08 | 0.13 | 0.008 | 0.28 | 0.33 | MA |
| S2_231152916 | 7 | 23,984,279 | 7.88E-07 | 0.08 | 0.06 | 0.011 | 0.30 | 0.34 | AllEnv |
| S8_152967929 | 8 | 157,645,450 | 1.92E-06 | 0.08 | 0.20 | 0.008 | 0.28 | 0.33 | MA |
| S8_152967952 | 8 | 157,645,473 | 1.08E-06 | 0.06 | 0.19 | 0.008 | 0.28 | 0.33 | MA |
| S8_152967966 | 8 | 157,645,487 | 2.64E-06 | 0.09 | 0.20 | 0.007 | 0.28 | 0.32 | MA |
| S10_144686998 | 10 | 145,399,603 | 1.08E-06 | 0.08 | 0.23 | 0.011 | 0.30 | 0.34 | AllEnv |

<sup>a</sup>SNP name is provided as “S” following by chromosome number and physical position in base pair from B73 Refgen\_v2. The genomic position of S2\_231152916 changed from chromosome 2 to 7 after uplifting from B73 RefGen\_v2 to RefGen\_v4.
<sup>b</sup>Genomic position (bp) of the SNP from B73 Refgen\_v4
<sup>c</sup>False discovery rate adjusted P-value
<sup>c</sup>False discovery rate adjusted P-value
<sup>e</sup> $R^2_{LR}$ likelihood ratio value of model without SNP
<sup>f</sup> $R^2_{LR-SNP}$ likelihood ratio value of model with SNP

In the Wisconsin Diversity panel, genome-wide LD decayed to background levels (*r*^2^ < 0.1) by 213 kb at the 90% percentile of the *r^2^* distribution (Figure S7). This percentile cutoff of the distribution was used to sample the large variance in LD structure presumably caused by rare variants (Wallace *et al*. 2014) and allow for the inclusion of potential distant regulatory elements as has been observed for cloned quantitative trait loci (QTL) in maize (Salvi *et al*. 2007; Studer *et al*. 2011). Therefore, the genomic search space to identify candidate genes was restricted to ± 200 kb window of the five peak SNPs that each tagged a unique genomic interval, resulting in the inclusion of 76 annotated genes (Table S8). Of these 76 genes, 57 are canonical protein coding genes, while the other 19 encode non-coding RNAs.

The strongest association signal was identified on chromosome 4 (Figure 1), with SNPs S4_30201436 and S4_30201599 comprising the peak associations with *g*_c_ in MA (*P*-value 2.68 × 10^-7^) and AllEnv (*P*-value 1.04 × 10^-6^), respectively. The peak SNP S4_30201599 resides within a microRNA locus (zma-MIR2275a) and is located 1,146 bp from a gene (*Zm00001d049479*) encoding an INCREASED SALT TOLERANCE1-LIKE1 (ISTL1) protein (Figure 2; Table S8). ISTL1 is the most probable candidate gene underlying this association signal because it is predicted to function together with the endosomal sorting complex required for transport machinery (ESCRT) and ATPase VACUOLAR PROTEIN SORTING4 (VPS4) in the formation of multivesicular bodies (MVBs) (Hill and Babst 2012; Buono *et al*. 2016).

**Figure 2.**
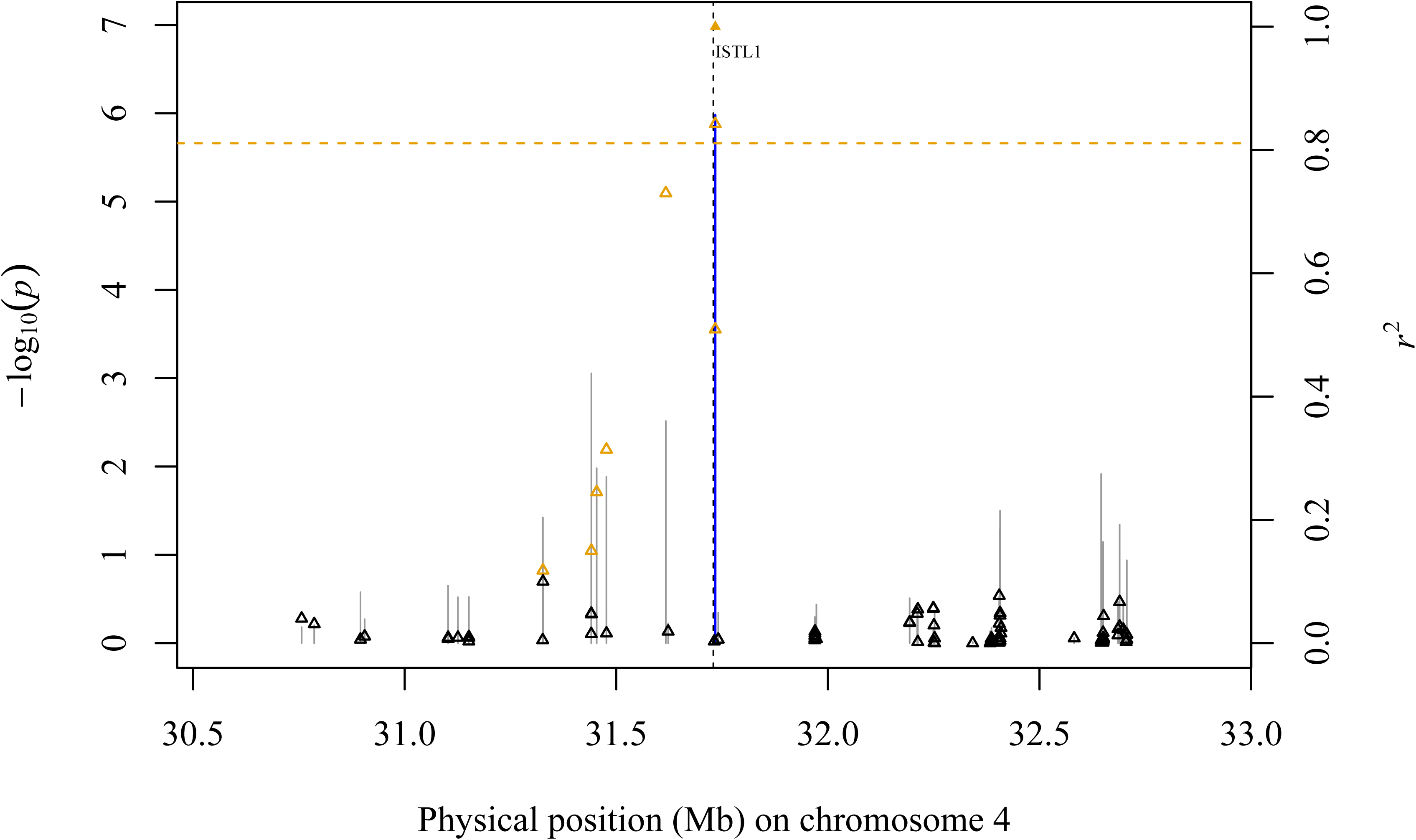
Association of SNP markers with adult maize leaf cuticular conductance (*g*_c_) across a genomic region on chromosome 4. Scatter plot of association results from a mixed linear model analysis of *g*_c_ conducted across all four environments (AllEnv) and linkage disequilibrium (LD) estimates (*r*^2^) for a genomic region that contains a gene encoding an INCREASED SALT TOLERANCE1-LIKE1 (ISTL1) protein (*Zm00001d049479*). The −log_10_ *P*-values of tested single-nucleotide polymorphisms (SNPs) are represented by vertical lines. Blue vertical lines are SNPs that are statistically significant at a false discovery rate (FDR) of 10%. The *r*^2^ values of each SNP relative to the peak SNP (indicated by a solid orange triangle) at 31,733,973 bp (B73 RefGen_v4) are indicated by triangles. Open orange triangles represent SNPs with *r*^2^ > 0.1 relative to the peak SNP. The least significant SNP at a genome-wide FDR of 10% is indicated by a dashed horizontal orange line. The black dashed vertical line indicates the genomic position of the ISTL1 protein.

The genomic region associated with *g*_c_ on chromosome 1 was detected in MA, with the signal peak defined by SNP S1_27055151534 (*P*-value 4.37 × 10^-7^) (Table 2). The peak SNP is located 8,190 bp from the nearest gene (*Zm00001d033835*), but this gene encodes an uncharacterized protein. Two genes adjacent to this one encode closely related SEC14 homologs, separated by about 50 kb (*Zm00001d033836*, ∼50 kb from peak SNP; and *Zm00001d033837*, ∼100 kb from peak SNP) (Figure S8; Table S8). SEC14 proteins in plants and other eukaryotes function in the transfer of phosphoinositides between different cellular membranes (Huang *et al*. 2016) and are noteworthy candidates for regulators of *g*_c_ because of their function in stimulating vesicle formation from the trans Golgi network in yeast (de Campos and Schaaf 2017), given the role of Golgi-dependent vesicle trafficking in cuticle biosynthesis in plants (McFarlane *et al*. 2014). Additional plausible candidates on chromosome 1 encode a homolog of ECERIFERUM7 (CER7) (*Zm00001d033842*) and a CYCLASE-ASSOCIATED PROTEIN (CAP) (*Zm00001d033830*) located (+) 179,198 bp and (-) 152,746 bp from the peak SNP, respectively (Figure S8). CER7, a 3′-to-5′ exoribonuclease family protein, is a core subunit of the RNA degrading exosome that positively regulates the expression of *CER3/WAX2/YRE*, a key wax biosynthesis gene in Arabidopsis (Kurata *et al*. 2003; Chen *et al*. 2003). CAP, a regulator of actin cytoskeleton dynamics, is highly conserved in plants, yeast, flies, and mammals (Balcer *et al*. 2003; Ono 2013).

An additional association with *g*_c_ was detected in MA on chromosome 8, defined by S8_152967952 as the peak SNP (*P*-value 1.92 x 10^-6^) along with two neighboring SNPs (S8_152967929 and S8_152967966). The closest gene (*Zm00001d011655*) is located 4,067 bp from the peak SNP, and it encodes a MITOGEN-ACTIVATED PROTEIN KINASE KINASE KINASE18. A gene encoding a GLY-ASP-SER-LEU ESTERASE/LIPASE (GDSL lipase) (*Zm00001d011661*) was identified 170,643 bp from the peak SNP (Figure S9; Table S8), and it was identified as the most plausible candidate in this region based on the known functions of GDSL lipases in lipid biosynthesis (Girard *et al*. 2012; Yeats *et al*. 2012b).

Finally, in AllEnv, *g*_c_ was also significantly associated with genomic regions on chromosomes 7 and 10 with peak SNPs S2_231152916 (*P*-value 7.88 x 10^-7^) and S10_144686998 (*P*-value 1.08 x 10^-6^), respectively. The peak SNP on chromosome 7 is located within a gene encoding CYTOKININ-N-GLUCOSYLTRANSFERASE1 (*Zm00001d019250*) (Figure S10; Table S8). The peak SNP on chromosome 10 is located within a gene that encodes β-D-XYLOSIDASE 4 (*Zm00001d026415*), a cell wall biosynthetic enzyme (Figure S11; Table S8). The latter is a plausible candidate given that the outer epidermal cell wall and cuticle are closely connected and polysaccharides can be found embedded in some parts of the cuticle (Yeats and Rose 2013). Therefore, sugar-or cell wall-modifying enzymes might directly or indirectly influence cuticle composition and permeability (Müller *et al*. 2013).

### Transcript abundance analysis of identified candidate genes

We analyzed the transcript abundance of the 57 candidate genes that encode a protein to help further prioritize which of them may be involved in the genetic control of *g*_c_. Given that the leaf cuticle is produced by epidermal cells, we analyzed LM-RNAseq data of epidermal cells from seven 2-cm intervals excised from the expanding leaf 8 of maize inbred B73 (Qiao *et al*. 2019), representing sequential stages in cuticle maturation (Bourgault *et al*. 2019). Of the 57 candidate genes, 24 were found to be expressed in at least one of the seven sampled intervals (Figure S12).

Of the seven promising candidate genes contributing to the genetic regulation of natural variation for *g*_c_ through potential influence on cuticle development, those encoding CAP, ISTL1 protein, GDSL lipase, β-D-XYLOSIDASE 4, and the two SEC14-like proteins were found to be epidermally expressed (Figure 3), while the homolog of CER7 was not declared to be expressed in the leaf epidermis based on its transcript abundance (Table S9). In general, transcript abundance increased for the ISTL1 protein and GDSL lipase along the leaf developmental gradient from the base (youngest, 2-4 cm) towards the tip (oldest, 20-22 cm) of the leaf, with the exception of a reduced transcript abundance for GDSL lipase in the 20-22 cm interval, at which time cuticle maturation is already completed (Bourgault *et al*. 2019). In contrast, CAP showed the opposite trend, with decreases in transcript abundance across the gradient from 2-4 cm to 20-22 cm intervals. The two genes encoding the SEC14-like proteins, *Zm00001d033836* and *Zm00001d033837*, had relatively constant transcript abundances from the base to the tip, with a slightly increased abundance at the 20-22 cm interval for *Zm00001d033837*. The gene encoding β-D-XYLOSIDASE 4 had a peak abundance at the 6-8 cm interval, followed by a progressive decrease in abundance towards the leaf tip. These observed transcript abundance patterns are compatible with roles in cuticle maturation.

**Figure 3.**
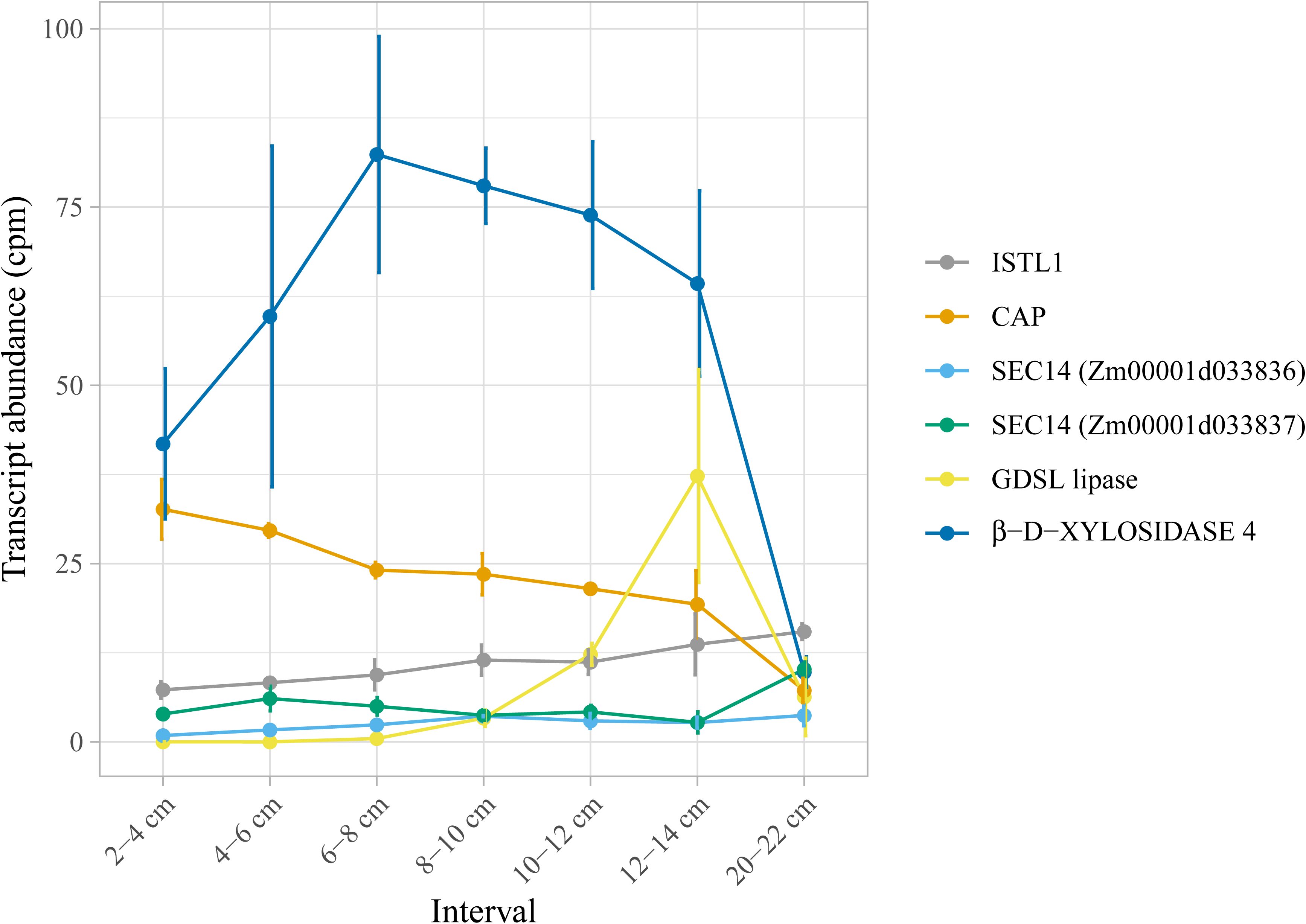
Transcript abundance of six candidate genes in maize leaf epidermal cells. Transcript abundance (counts per million, cpm) of each gene is plotted against the developmental gradient (six intervals from 2–14 cm, and one interval from 20–22 cm) of expanding adult leaf 8 from maize inbred B73. Error bars represent the standard deviation.

### Whole-genome prediction of *g*_c_

Whole-genome prediction (WGP) was performed with the complete marker data set of 235,004 SNPs to assess the feasibility of implementing genomic selection for the genetic improvement of *g*_c_ in maize breeding populations. Predictive ability was evaluated on *g*_c_ from MA, SD, and AllEnv in a scheme that used each of the three as a training set. The overall predictive ability was 0.48 across all of the nine training-prediction combinations, with individual predictive abilities ranging from 0.40 to 0.54 (Figure 4; Table S10). Not unexpectedly, the highest average predictive ability (0.52) was achieved when AllEnv was the training set, followed by MA (0.46) and SD (0.45). Indicative of contrasting environmental conditions between field locations, MA and SD showed lower predictive abilities for each other compared to when MA or SD was used to predict itself. However, AllEnv as the training set resulted in more similar prediction abilities for *g*_c_ in MA (0.53) and SD (0.49).

**Figure 4.**
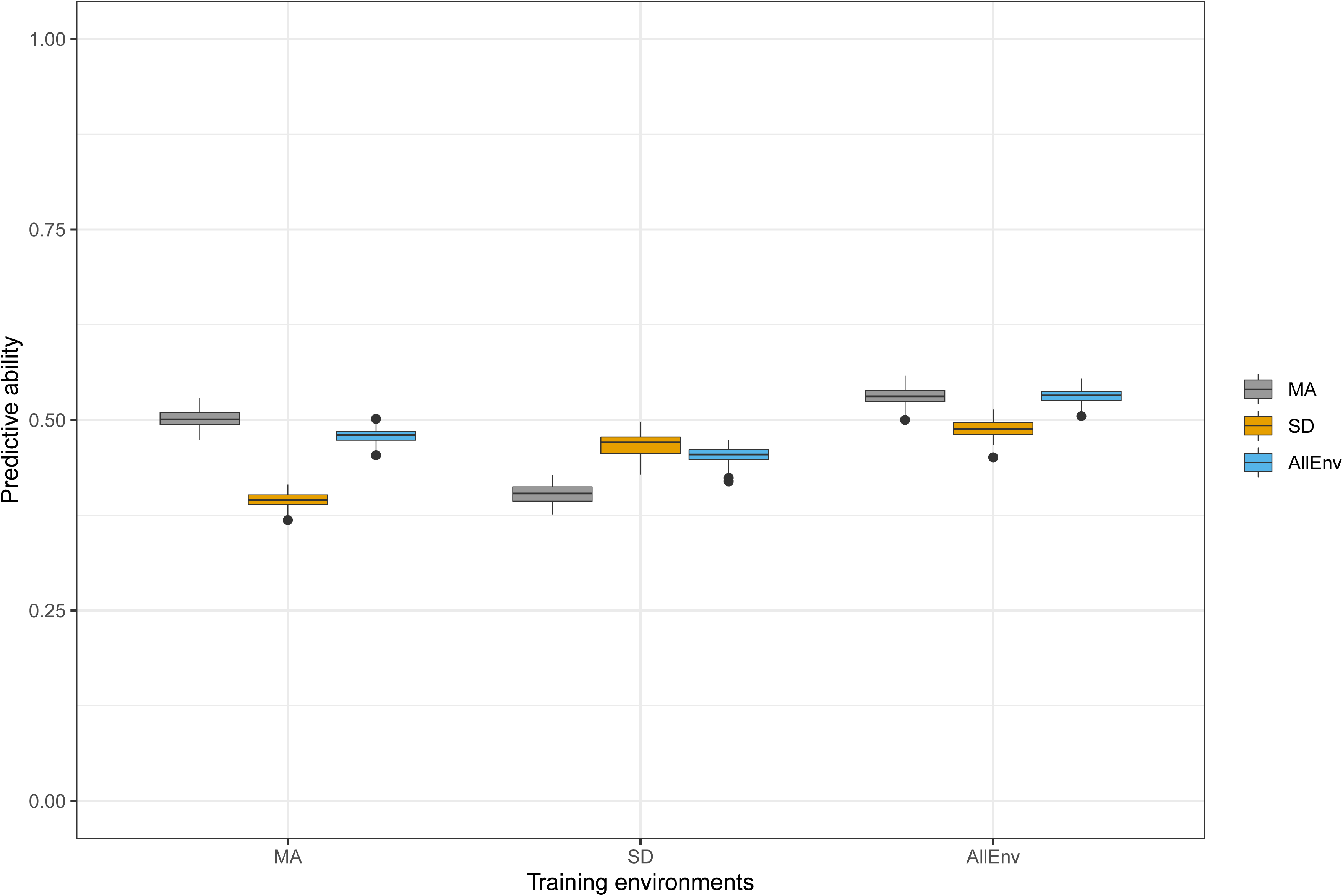
Predictive abilities of whole-genome prediction for adult maize leaf cuticular conductance (*g*_c_) in locations (Maricopa, MA; and San Diego, SD) and across all four environments (AllEnv) in a scheme that used each of the three as a training set. Grey, orange and blue represent the predictive abilities for *g*_c_ in MA, SD, and AllEnv, respectively.

## Discussion

The cuticle, a crucial barrier to plant water loss (Shepherd and Wynne Griffiths 2006; Xue *et al*. 2017), has been widely studied using single gene loss- and gain-of-function mutants, and interspecies crosses (Kosma and Jenks 2007; Yeats *et al*. 2012a; Beisson *et al*. 2012; Yeats and Rose 2013). However, no prior study has explored natural variation in a single species to elucidate the genetic control of cuticle function as a water barrier on a genome-wide scale. In this study, we focused on leaf cuticular conductance (*g*_c_), a trait of potential agronomic value related to drought tolerance. To that end, we evaluated the phenotypic variation of *g*_c_ using detached adult leaves in a maize diversity panel in two climatically contrasting field locations (Table S1) and conducted a GWAS and WGP to explore the genetic architecture and the potential genetic gains that could be expected under selection in a breeding program for *g*_c_.

Analysis was performed to evaluate the repeatability of *g*_c_ in and across locations, as well as how *g*_c_ related to flowering time (DTA). Moderately strong correlations (*r*) of *g*_c_ measurements were observed between years in a location, while relatively weaker correlations existed between locations (Figure 5). Given the lower relative humidity and higher solar radiation and air temperature of MA compared to SD during maize growing seasons (Table S1), the weaker correlative patterns between locations imply that *g*_c_ is moderately responsive to environmental conditions. Noticeably, weak to moderately strong negative correlations (MA16, −0.33; MA17, −0.50; SD16, −0.14; SD17, −0.20) existed between *g*_c_ and flowering time collected from the same year in each location. These findings suggest low to moderate levels of confounding between *g*_c_ and maturity depending on the environment, with the prolonged exposure of the developing cuticle of plants to more extreme weather variables possibly explaining the stronger correlations in the MA field location. Even though these results implicated the need to assess for the confounding effect of flowering time when conducting GWAS, the ∼2-fold range in *g*_c_ variation, calculated as the ratio between the maximum and minimum BLUP values, in (MA and SD) and across (AllEnv) locations, and the moderately high heritabilities of *g*_c_ (Table 1) indicate that this phenotype should be very amenable to genetic dissection and prediction in maize.

We conducted a GWAS that resulted in the identification of five genomic regions associated with *g*_c_ in the Wisconsin Diversity panel. Of these five genomic regions, two were detected only in MA (chromosomes 1 and 8), two in AllEnv (chromosomes 7 and 10), and one (chromosome 4) was detected in both MA and AllEnv (Figure 1; Table 2). Notably, no significant associations at the 10% FDR level were detected only in SD, implying that G×E exists for *g*_c_ as shown by the between-location correlation analysis (Figure S5). This is also further supported by comparing the ratio of genetic variance to G×E variance, which was 3.95, 2.57, and 1.81 for SD, MA, and AllEnv, respectively (Table S5). All together, these observations are consistent with those of previous studies showing that both the content and composition of cuticular waxes can be influenced by environmental factors such as ultraviolet (UV) radiation, temperature, humidity, and drought stress (Shepherd and Wynne Griffiths 2006; Xue *et al*. 2017). It has been reported that drought stress and low humidity increased wax loads in many plant species (Sutter and Langhans 1982; Shepherd and Wynne Griffiths 2006; Koch *et al*. 2006; Kosma and Jenks 2007), but a different combination of UV intensity and temperature resulted in more complex patterns of wax products (Baker 1974; Riederer and Schneider 1990; Shepherd *et al*. 1997). Although it is not straightforward to explain how each environmental factor affected *g*_c_ through wax biosynthesis and deposition, the differences in GWAS results in MA and SD suggests that other environment-specific factors in MA besides the high temperature, elevated UV radiation, and low humidity (Supplemental Table S1) might be inducing compositional features that affect water loss relative to plants in SD.

As the major barrier of water vapor diffusion from the intercellular air space to the atmosphere when stomata are closed, the composition and structure of the cuticle can affect water evaporation resistance, thus causing variation in *g*_c_. The candidate genes we propose for *g*_c_ via GWAS encode proteins with functions that can be related to cuticle formation.

Three of the candidate genes implicated by our GWAS in control of *g*_c_ encode proteins with likely functions in membrane trafficking, a process needed for deposition of cuticle lipids at the cell surface as well as for delivery of lipid transport proteins to the plasma membrane (McFarlane *et al*. 2014). Two adjacent genes, likely resulting from a recent tandem duplication, encode closely related SEC14 homologs. These belong to a subset of SEC14 homologs in maize lacking the GOLD (Golgi dynamics) and nodulin domains present in SEC14s that have been functionally characterized in plants (Huang *et al*. 2016; de Campos and Schaaf 2017). Nevertheless, SEC14 domains are well known to function in movement of phosphoinositides (signaling lipids) between cellular membranes; in yeast this function stimulates vesicle formation from the trans-Golgi network (de Campos and Schaaf 2017). Thus, the maize SEC14 proteins we identified may function in vesicle trafficking as well.

Another membrane-trafficking-related candidate encodes an IST1 family protein closely related to Arabidopsis ISTL1 (Buono *et al*. 2016). Yeast IST1 functions with the ATPase VPS4 to regulate the assembly and function of ESCRTIII complexes, which facilitate the formation of vesicles inside MVBs (Hill and Babst 2012). MVBs are late endosomal compartments that function in degradation of proteins retrieved via endocytosis from the plasma membrane. A function for ISTL1 proteins in degradation of plasma membrane proteins and formation of MVBs in Arabidopsis has been demonstrated (Buono *et al*. 2016). Thus, maize ISTL1 could function in an endosomal pathway critical for maintenance of plasma membrane-associated proteins that deliver cuticle lipids to the extracellular environment such as ABCG transporters and lipid transfer proteins. Moreover, MVB-like organelles have been implicated as the source of extracellular vesicles that deliver certain proteins and small RNAs to the extracellular environment in plants (Rutter and Innes 2018). This suggests the additional possibility that maize ISTL1 could impact *g*_c_ because of a role in the formation of MVB vesicles containing cuticle lipids or GDSL lipases required for cutin polymerization, which may be released into the extracellular environment via fusion of MVBs with the plasma membrane. Confirmation of a role for MVBs in cuticle formation would reveal new aspects of membrane trafficking supporting cuticle formation.

CYCLASE-ASSOCIATED PROTEIN1, encoded by the candidate gene on chromosome 1 near peak SNP S1_270551534, has not been associated with cuticle development in previous studies. CAPs are actin monomer-binding proteins that regulate actin dynamics (Qualmann *et al*. 2000; Hubberstey and Mottillo 2002). Functions for CAPs in actin regulation and actin-dependent processes in Arabidopsis have been demonstrated through genetic and biochemical studies (Barrero *et al*. 2002; Chaudhry *et al*. 2007; Deeks *et al*. 2007). Actin dynamics have been implicated in multiple aspects of membrane trafficking in eukaryotic cells including vesicle formation and vesicle movement (Lanzetti 2007). Thus, CAP could impact *g*_c_ in maize via regulation of membrane trafficking events supporting cuticle formation such as Golgi-mediated secretion (McFarlane *et al*. 2014; Luo *et al*. 2019) and/or endocytosis.

Two of the candidate genes we identified have predicted functions in cuticle formation. One of these is CER7, which functions in Arabidopsis to regulate the biosynthesis of alkanes, one of the major classes of cuticle waxes in this species (Jenks *et al*. 2002) and also in maize adult leaves(Bourgault *et al*. 2019). CER7 encodes an exosomal 3′-to-5′ exoribonuclease that regulates the expression level of *CER3* (Hooker *et al*. 2007). The CER3 protein acts together with CER1 to catalyze the formation of alkanes (Bernard *et al*. 2012). The other candidate with a predicted function in cuticle formation encodes a GDSL lipase (*Zm00001d011661*). GDSL lipases catalyze the hydrolysis of mono-, di- and triglycerols and release free fatty acids and alcohols (Angkawidjaja and Kanaya 2006). Although GDSL lipases are involved in a large number of biological processes in plants, their function in the biosynthesis of cutin is well-recognized(Takahashi *et al*. 2010; Girard *et al*. 2012; Yeats *et al*. 2012b). Moreover, a homolog of *Zm00001d011661* in Arabidopsis, *AT3G16370*, was down-regulated along with other cuticle associated genes in a transgenic DESPERADO/ABCG11 silenced line with changes in levels of cutin monomers (Panikashvili *et al*. 2010), whereas its homolog in *Citrus clementina* was reported as the top differentially expressed gene in the fruit epidermis(Matas *et al*. 2010).

The final candidate gene identified encodes the cell wall biosynthesis enzyme β-D-XYLOSIDASE 4 (*Zm00001d026415*). Sometimes the cuticle is described as a specialized lipidic modification of the cell wall, and polysaccharides are known to be deposited in some regions of the cuticle (Yeats and Rose 2013). Transmission electron microscopy and non-destructive polarization modulation-infrared reflection-absorption spectroscopy have challenged the long-standing notion that polysaccharide compounds, mostly pectin, are confined to the cuticular layer of the cuticle, adjacent to the cell wall. Other polysaccharides, hemicelluloses such as xylan and xyloglucan, were found close to the surface of the cuticle, within the cuticle proper of several species (Guzmán *et al*. 2014; Hama *et al*. 2019). Therefore, cell wall-modifying enzymes could have an influence on the polysaccharide content of the cuticle, or might impact cuticle composition or organization by affecting the transit of cuticle lipids across the wall to the cuticle (Müller *et al*. 2013).

Genes encoding CAP, ISTL1, GDSL lipase, BETA-D-XYLOSIDASE 4, and both SEC14 homologs were declared expressed in epidermal cells of developing maize leaves (Figure 3), where the cuticle matures, indicating that these genes are involved in metabolic and other cellular processes in maize leaf epidermal cells and possibly associated with cuticle development. The abundance of the CER7 homolog transcript, a core subunit of the exosome that regulates wax biosynthesis (Hooker *et al*. 2007), was too low to be declared expressed, with cpm values ranging from 0.033 to 0.137 in 6 out of the 20 samples. Similarly, very low abundances of this transcript were observed by RNAseq in a number of different tissues from maize inbred B73 (Stelpflug *et al*. 2016). The increasing/decreasing trends in transcript abundance along the developmental gradient (Figure 3) are consistent with regulatory roles in cuticle maturation; however, these trends are equally consistent with roles in other developmental changes taking place at the same time. Additional work will be needed to definitively link changes in expression levels of these genes with variation in g_c_.

In our GWAS results, even after controlling for the confounding impact of flowering time, each genomic region significantly associated with *g*_c_ explained 4-6% (*R^2^_LR_*_-SNP_ - *R^2^_LR_*) of the variation in the panel (Table 2), thus only accounting for a minor fraction of the estimated heritability. If the genetic architecture of *g*_c_ is predominated by a large number of loci with small allelic effect sizes, then it is possible that our association population was not of a sample size sufficient to offer the statistical power needed to detect functional variants with small effects (Long and Langley 1999). With only a total of 235,004 SNPs with common minor allele frequencies (5% or greater) scored on the diversity panel, the “missing” heritability could be also partially attributed to not having a density of SNP markers required to achieve strong LD (*r^2^* > 0.80) with unscored causal variants that are potentially rare in frequency (Myles *et al*. 2009; Buckler *et al*. 2009). Therefore, increasing SNP marker density and the mapping population size are likely important for explaining the missing heritability of *g*_c_, which is possibly controlled by large numbers of small effect alleles in maize considering the complex cuticle biosynthesis network and transport mechanisms (Yeats and Rose 2013), as well as factors that affect leaf water trafficking.

In general, the predictive abilities of WGP were moderately high for the tested nine training-prediction combinations (Figure 4; Table S10), indicating that genetic gain for *g*_c_ could be accelerated with WGP in maize breeding programs. Presumably due to influence of contrasting environmental conditions on the phenotypic data collected at the two field locations (Figure S5; Tables S1 and S4), the prediction abilities were lowest when the MA and SD training sets predicted *g*_c_ each other (Figure 4; Table S10). However, AllEnv showed similar predictive abilities for MA and SD as for themselves, because AllEnv contained phenotypic information from both locations. Given the range in phenotypic variation observed for *g*_c_ across locations, it is highly recommended that phenotyping and selection occur in the specific target breeding environments, especially if the focus is on hot, arid environments such as MA. Furthermore, if *g*_c_ does ultimately have a polygenic genetic basis, genomic selection would be the optimal breeding strategy for *g*_c_ and inevitably drought tolerance in maize breeding populations rather than a marker-assisted selection approach designed for only a few loci with large effects (Meuwissen *et al*. 2001; Lorenz *et al*. 2011).

## Acknowledgements

We especially thank Aaron Waybright, Christine Caron, Angel Mendoza, Albert Nguyen, Indira Oueralta Castillo, Anastasia Zagordo, and the BICD 101 students at UCSD in the summer of 2016 and 2017, for collecting phenotypic data. We thank Mark Millard and Candice Gardner of the USDA-ARS North Central Regional Plant Introduction Station (NCRPIS) in Ames, Iowa, and Candice Hirsch at the University of Minnesota for providing seed of the Wisconsin Diversity panel. We also thank Bill Luckett, Andrew French, Kelly Thorp, Alison Thompson, John Dyer and others at the U.S. Arid-Land Agricultural Research Center in Maricopa, AZ, for their assistance with planting and providing a facility for phenotyping. Additionally, we thank Clint Jones, Greg Main, Russell Noon, Rick Ward and others at the University of Arizona, Maricopa Agricultural Center in Maricopa, AZ, for the management of the Arizona field trials. This research was supported by the National Science Foundation IOS1444507.

## Supplemental information

**Supplemental Figure S1.**

Diffusion conductance for adult maize leaves transferred from daylight into the dark. Adult maize leaves were excised from field-grown plants in the same manner routinely used for leaf cuticular conductance (*g*_c_) phenotyping, and transferred to a dark room where their cut bases were submerged in water. Diffusion conductance was measured immediately (time zero), and at successive 30 min intervals, to determine the amount of time needed to reach minimum conductance values in the dark. Panels A-I display results for each genotype tested (indicated at the top of the panel) with each line representing a different plant (1-3 plants tested for most genotypes and 7 plants for B73, a reference standard harvested each sampling day). All analyzed inbred lines except for B73, which was the reference standard, were in the top 15% for *g*_c_ across all four environments (AllEnv).

**Supplemental Figure S2.**

The degree of relationship (Pearson’s correlation, *r*) between the plot-level averages of leaf surface area (mm^2^) and leaf dry weight (g) collected from the same adult maize leaves used for phenotyping cuticular conductance (*g*_c_) in Maricopa and San Diego in 2017.

**Supplemental Figure S3.**

Population structure analysis of the Wisconsin Diversity panel. (A) Principal component analysis (PCA) plot colored by group classification of lines [Flint, Iodent, Non-Stiff Stalk (NSS), Stiff Stalk Synthetic (SSS), Popcorn, Sweet corn, Tropical, Unknown (i.e., genotyped but ambiguous group), and Not available (i.e., not genotyped)] following that of Hansey et al. (2011). (B) Distruct plot of fastSTRUCTURE subpopulation membership with K = 6. (C) PCA plot colored according to the fastSTRUCTURE subpopulation membership with K=6. Individuals with assignment values (Q) less than 50% were classified as admixed. Populations 1, 2, and 4 predominantly contained lines belonging to the NSS group; population 3 mostly had lines from the popcorn and sweet corn groups; population 5 mainly included lines from the tropical group; and population 6 primarily had lines from the SSS group.

**Supplemental Figure S4.**

Principal component analysis (PCA) of transcript abundance for the 21 samples collected from leaf 8 of maize inbred B73 in the LM-RNAseq analysis. The first two principal components (PCs) correspond to developmental stage (PC1) and tissue type (PC2). Each point corresponds to an RNAseq sample. From left to right, each color corresponds to a specific developmental stage (youngest to oldest part of the leaf). From bottom to top, circles represent epidermal tissues, and triangles represent internal tissues. The orange dot and triangle points that are encompassed by a black circle were considered an outlier for stage 2 (interval 4 - 6 cm), thus these samples were removed.

**Supplemental Figure S5.**

Correlation matrix for best linear unbiased predictors (BLUPs) of maize adult leaf cuticular conductance (*g*_c_; g·h^-1^·g^-1^) and flowering time (FT; days to anthesis) from each of four single environments [Maricopa (MA) and San Diego (SD) in 2016 and 2017; MA16, SD16, MA17, SD17], and the first three principal components calculated from the SNP genotype matrix. Pearson’s correlation coefficients (*r*) are presented in the upper right triangle, while the corresponding *P*-values for the significance of associations (α = 0.05) are displayed below the diagonal.

**Supplemental Figure S6.**

Genome-wide association study (GWAS) of adult maize leaf cuticular conductance (*g*_c_) conducted in locations (Maricopa, MA; and San Diego, SD) and across all four environments (AllEnv). The −log_10_ *P*-value of each SNP tested in a mixed linear model analysis of *g*_c_ is plotted as a point against its physical position (B73 RefGen_v4) for the 10 chromosomes of maize. The least significant single-nucleotide polymorphism (SNP) at a genome-wide false discovery rate of 10% is indicated by a dashed horizontal black line.

**Supplemental Figure S7.**

Linkage disequilibrium (LD) estimates in the Wisconsin Diversity panel. Mean decay of LD measured as pairwise *r*^2^ from the 235,004 SNP markers over physical distance. Black lines indicate the distribution of SNP markers at different percentile cutoffs. The grey horizontal line indicates *r*^2^ of 0.1.

**Supplemental Figure S8.**

Association of SNP markers with adult maize leaf cuticular conductance (*g*_c_) across a genomic region on chromosome 1. Scatter plot of association results from a mixed linear model analysis of *g*_c_ conducted in Maricopa (MA) and linkage disequilibrium (LD) estimates (*r*^2^) for a genomic region that contains genes encoding a CYCLASE-ASSOCIATED PROTEIN1 (CAP, *Zm00001d033830*), two SEC14-like proteins [SEC14 (3836), *Zm00001d033836*; SEC14 (3837), *Zm00001d033837*], and a homolog of ECERIFERUM7 (CER7, *Zm00001d033842*). The −log_10_ *P*-values of tested single-nucleotide polymorphisms (SNPs) are represented by vertical lines. Blue vertical lines are SNPs that are statistically significant at a false discovery rate (FDR) of 10%. The *r*^2^ values of each SNP relative to the peak SNP (indicated by a solid orange triangle) at 275,318,146 bp (B73 RefGen_v4) are indicated by triangles. Open orange triangles represent SNPs with *r*^2^ > 0.1 relative to the peak SNP. The least significant SNP at a genome-wide FDR of 10% is indicated by a dashed horizontal orange line. The black dashed vertical line indicates the genomic positions of the CAP, two SEC14-like proteins, and a CER7 homolog.

**Supplemental Figure S9.**

Association of SNP markers with adult maize leaf cuticular conductance (*g*_c_) across a genomic region on chromosome 8. Scatter plot of association results from a mixed linear model analysis of *g*_c_ conducted in Maricopa (MA) and linkage disequilibrium (LD) estimates (*r*^2^) for a genomic region that contains a gene encoding a protein of the GLY-ASP-SER-LEU ESTERASE/LIPASE (GDSL lipase, *Zm00001d011661*) on chromosome 8. The −log_10_ *P*-values of tested single-nucleotide polymorphisms (SNPs) are represented by vertical lines. Blue vertical lines are SNPs that are statistically significant at a false discovery rate (FDR) of 10%. The *r*^2^ values of each SNP relative to the peak SNP (indicated by a solid orange triangle) at 157,645,473 bp (B73 RefGen_v4) are indicated by triangles. Open orange triangles represent SNPs with *r*^2^ > 0.1 relative to the peak SNP. The least significant SNP at a genome-wide FDR of 10% is indicated by a dashed horizontal orange line. The black dashed vertical line indicates the genomic position of GDSL lipase.

**Supplemental Figure S10.**

Association of SNP markers with adult maize leaf cuticular conductance (*g*_c_) across a genomic region on chromosome 7. Scatter plot of association results from a mixed linear model analysis of *g*_c_ conducted across all four environments (AllEnv) and linkage disequilibrium (LD) estimates (*r*^2^) for a genomic region that contains the peak SNP. The - log_10_ *P*-values of tested single-nucleotide polymorphisms (SNPs) are represented by vertical lines. Blue vertical lines are SNPs that are statistically significant at a false discovery rate (FDR) of 10%. The *r*^2^ values of each SNP relative to the peak SNP (indicated by a solid orange triangle) at 23,984,279 bp (B73 RefGen_v4) are indicated by triangles. Open orange triangles represent SNPs with *r*^2^ > 0.1 relative to the peak SNP. The least significant SNP at a genome-wide FDR of 10% is indicated by a dashed horizontal orange line.

**Supplemental Figure S11.**

Association of SNP markers with adult maize leaf cuticular conductance (*g*_c_) across a genomic region on chromosome 10. Scatter plot of association results from a mixed linear model analysis of *g*_c_ conducted across all four environments (AllEnv) and linkage disequilibrium (LD) estimates (*r*^2^) for a genomic region that contains a gene encoding a β-D-XYLOSIDASE 4 (*Zm00001d026415*). The −log_10_ *P*-values of tested single-nucleotide polymorphisms (SNPs) are represented by vertical lines. Blue vertical lines are SNPs that are statistically significant at a false discovery rate (FDR) of 10%. The *r*^2^ values of each SNP relative to the peak SNP (indicated by a solid orange triangle) at 145,399,603 bp (B73 RefGen_v4) are indicated by triangles. Open orange triangles represent SNPs with *r*^2^ > 0.1 relative to the peak SNP. The least significant SNP at a genome-wide FDR of 10% is indicated by a dashed horizontal orange line. The black dashed vertical line indicates the genomic position of β-D-XYLOSIDASE 4.

**Supplemental Figure S12.**

Heatmap of average log_2_ transformed transcript abundance (counts per million; cpm) for the 24 candidate genes declared to be expressed in epidermal cells of expanding adult leaf 8 of maize inbred B73. A logarithmic scale was used due to the large range of cpm values across genes, and a constant with a value of 1 was added to all cpm values (cpm + 1) to prevent the generation of negative values after log_2_ transformation. Candidate genes are grouped by the five genomic regions associated with *g*_c_, which is indicated by the peak SNP of each genomic region.

**Supplemental Table S1.**

Summary of weather conditions for each field experiment conducted in Maricopa (MA) and San Diego (SD) in 2016 and 2017 (MA16, SD16, MA17 and SD17) from the day of planting to the last day of phenotyping leaf cuticular conductance (*g*_c_).

**Supplemental Table S2.**

Plot-level averages of leaf dry weight and leaf surface area for 451 maize inbred lines from Maricopa and San Diego in 2017 (MA17 and SD17). For each inbred line, an average of each trait was calculated based on all measured leaves from each plot.

**Supplemental Table S3.**

Best fitted models used to calculate best linear unbiased predictors (BLUPs) for adult maize leaf cuticular conductance (*g*_c_) and flowering time (days to anthesis) for each of four single environments [Maricopa (MA) and San Diego (SD) in 2016 and 2017; MA16, SD16, MA17, SD17], in a location (MA and SD), and across all four environments (AllEnv) according to a likelihood ratio test (α = 0.05). The star (*) indicates that a random effect term was retained in the mixed linear model, whereas the x indicates that a fitted random effect term was not significant and removed from the mixed linear model. The hyphen (-) indicates that a random effect term was not fitted in a mixed linear model.

**Supplemental Table S4.**

Best linear unbiased predictors (BLUPs) of adult maize leaf cuticular conductance (*g*c; g·h^-1^·g^-1^) and flowering time (FT; days to anthesis) for 451 maize inbred lines from each of four single environments [Maricopa (MA) and San Diego (SD) in 2016 and 2017; MA16, SD16, MA17, SD17], in a location (MA and SD), and across all four environments (AllEnv).

**Supplemental Table S5.**

Variance component estimates from the fitted full mixed linear model used to calculate heritability on a line-mean basis for leaf cuticular conductance (*g*_c_) and flowering time (days to anthesis) in a location (Maricopa, MA; and San Diego, SD) and across all four environments (AllEnv).

**Supplemental Table S6.**

Population structure analysis of the Wisconsin Diversity panel. Included in the table are the first three principal components (PCs) calculated from the SNP genotype matrix (PC1, PC2, and PC3), assignment value (Q) for each subpopulation (subpopulations 1 to 6), group classification of lines following assignments of Hansey et al. (2011), subpopulation with the highest assignment value (K), greatest Q assignment value, and the subpopulation (subpop. 1, 2, 3, 4, 5, 6, or admixed) to which each line was assigned (Category) in our study.

**Supplemental Table S7.**

Averages and ranges for best linear unbiased predictors (BLUPs) of flowering time (days to anthesis) in a location (Maricopa, MA; and San Diego, SD) and across all four environments (AllEnv), and estimated heritabilities on a line-mean basis.

**Supplemental Table S8.**

Genomic information (RefGen_v4) for the 76 candidate genes within ± 200 kb of the peak SNPs identified in the genome-wide association study of adult maize leaf cuticular conductance (*g*_c_) in a location (Maricopa, MA; San Diego, SD) and across all four environments (AllEnv).

**Supplemental Table S9.**

Transcript abundance levels as counts per million (cpm) for the 76 candidate genes within ± 200 kb of the peak SNPs identified in the genome-wide association study of adult maize leaf cuticular conductance (*g*_c_).

**Supplemental Table S10.**

Predictive abilities of whole-genome prediction models for adult maize leaf cuticular conductance (*g*_c_) in locations (Maricopa, MA; and San Diego, SD) and across all four environments (AllEnv).

